# Peripheral blood mononuclear cells exhibit increased mitochondrial respiration after adjuvant chemo- and radiotherapy for early breast cancer

**DOI:** 10.1101/2022.12.22.521564

**Authors:** Ida Bager Christensen, Lucas Ribas, Kristian Buch-Larsen, Djordje Marina, Steen Larsen, Peter Schwarz, Flemming Dela, Linn Gillberg

**Affiliations:** Department of Biomedical Sciences, University of Copenhagen, Copenhagen, Denmark; Department of Endocrinology and Metabolism, Rigshospitalet, Copenhagen, Denmark; Clinical Research Centre, Medical University of Bialystok, Bialystok, Poland; Faculty of Health and Medical Sciences, University of Copenhagen, Copenhagen, Denmark; Department of Geriatrics, Bispebjerg University Hospital, Copenhagen, Denmark

**Keywords:** Breast cancer, mitochondria, chemotherapy, radiotherapy, energy metabolism, high-resolution respirometry

## Abstract

**Background:** Adjuvant chemo- and radiotherapy cause cellular damage not only to cancerous but also to healthy dividing cells. Antineoplastic treatments have been shown to cause mitochondrial respiratory dysfunction in non-tumorous tissues, but the effects on circulating human peripheral blood mononuclear cells (PBMCs) remain unknown.

**Aim:** We aimed to identify changes in mitochondrial respiration of PBMCs after adjuvant chemo- and radiotherapy in postmenopausal early breast cancer (EBC) patients and relate these to metabolic parameters of the patients.

**Methods:** Twenty-three postmenopausal women diagnosed with EBC were examined before and shortly after chemotherapy treatment often administered in combination with radiotherapy (n=18). Respiration (O_2_ flux per million PBMCs) was assessed by high-resolution respirometry of intact and permeabilized PBMCs. Clinical metabolic characteristics were furthermore assessed.

**Results:** Respiration of intact and permeabilized PBMCs from EBC patients was significantly increased after adjuvant chemo- and radiotherapy (*p*=6×10^−5^ and *p*=1×10^−7^, respectively). The oxygen flux attributed to specific mitochondrial complexes and respiratory states increased by 17-43% compared to before therapy commencement. Leukocyte counts (*p*=1×10^−4^), hemoglobin levels (*p*=0.0003), and HDL cholesterol (*p*=0.003) decreased while triglyceride (*p*=0.01) and LDL levels (*p*=0.02) increased after treatment suggesting a worsened metabolic state. None of the metabolic parameters correlated significantly with PBMC respiration.

**Conclusion:** This study shows that mitochondrial respiration in circulating PBMCs is significantly increased after adjuvant chemo- and radiotherapy in postmenopausal EBC patients. The increase might be explained by a shift in PBMC subpopulation proportions towards cells relying on oxidative phosphorylation rather than glycolysis or a generally increased mitochondrial content in PBMCs. Both parameters might be influenced by therapy-induced changes to the bone marrow or vascular microenvironment wherein PBMCs differentiate and reside.

## Introduction

Breast cancer was the most frequently diagnosed malignancy in 2020 with 2.3 million new cases globally^1^. Fortunately, breast cancer patients have one of the highest five-year relative survival rates of 90 %^2^, attributed to mammography screening programs leading to earlier detection as well as improved treatment options^3,4^. Breast cancer therapy is often multimodal comprising surgery and adjuvant treatments including chemotherapy, radiotherapy, and/or anti-estrogen treatment combined to maximize therapeutic efficacy and minimize cancer recurrence, drug resistance, and undesired side effects^3,5^. However, these efficient and comprehensive treatment regimens of breast cancer patients are associated with whole-body metabolic derangements and endocrine side effects^6–10^. Among the documented metabolic changes are weight gain^11,12^, increased fasting insulin and HbA1c^13^ alongside decreased peripheral insulin sensitivity^14^ and increased risk of type 2 diabetes^15,16^ and cardiovascular disease^17–19^. Changes in lipid profile e.g., decreased HDL cholesterol and increased triglyceride levels are also described in breast cancer patients after chemotherapy^9,14^. Thus, metabolic deterioration in breast cancer patients after antineoplastic therapy is well established but underlying mechanisms have yet to be fully elucidated.

Cytotoxicity of chemotherapeutic drugs to malignant cells is achieved by interference with growth kinetics, e.g., via direct DNA alkylation damage^20^, intercalation of drug molecules into nucleic acids^21^, microtubule disruption^22^, or enzymatic interference^23^. Radiotherapy deposits high-energy ionizing radiation localized onto malignant tissues inducing DNA damage and cell death^24^. However, antineoplastic treatments and local radiotherapy are not restricted to target cancer cells but affect the DNA and cellular functions in healthy cells as well^25,26^. For instance, chemotherapy has been shown to induce mitochondrial dysfunction^27–31^. Repeated administration of the chemotherapeutic anthracycline doxorubicin in mice impairs mitochondrial respiration in skeletal myocytes^27^ and causes e.g. deletion and oxidation of mitochondrial DNA and respiratory chain defects such as reduced activity of complexes of the electron transport chain in murine cardiomyocytes^28–31^. The effect of different antineoplastic drugs on mitochondrial oxygen consumption has been investigated by direct incubation of cells *in vitro*^32,33^. In one study, the alkylating agent cisplatin caused no distinct impairment of oxygen consumption whereas the anthracyclines doxorubicin and daunorubicin did inhibit mitochondrial respiration in leukemia and lymphoma cells.^32^. Ionizing radiotherapy is known to damage nuclear DNA but a direct effect on complex I, II, and III in the electron transport chain and ATP synthase activity has also been described, suggesting that radiation can also affect oxidative phosphorylation (OXPHOS)^34^. Interestingly, a recent study on skeletal myocytes from early (non-metastatic) breast cancer (EBC) patients before and after receiving (neo)adjuvant chemotherapy reports unchanged mitochondrial respiratory capacity but significantly reduced mitochondrial biogenesis and exacerbated H_2_O_2_ production^35^. To summarize, antineoplastic agents have been shown to affect mitochondrial function including respiratory capacity in various cell types exceeding cancer cells.

Mitochondrial respiration of peripheral blood mononuclear cells (PBMCs) has attracted attention since increasing number of studies indicate that the bioenergetic status of PBMCs is affected in several diseases^36–42^. Disturbed PBMC respiration has been reported for patients with septic shock^37^, type 2 diabetes^38^, end-stage renal disease^40^, age-related fatigue^41^, chronic fatigue syndrome^42^, and major depression disorders^39^ where respiratory impairment of PBMCs in the diseased state are typically reported. Circulating PBMCs are exposed to vascular microenvironments throughout the body, including systemic medical treatments, such as statins, which have been associated with increased mitochondrial respiration in PBMCs^43^. However, changes to the bioenergetic status of PBMCs from EBC patients after receiving adjuvant chemo- and radiotherapy remains unknown.

As circulating PBMCs have a substantial role in supporting immune function, knowledge of how systemic chemotherapy and localized radiotherapy affect important non-tumorous immune cells is of great importance. We hypothesized that systemic chemotherapy treatment affects the respiratory function of PBMCs. Thus, we aimed to investigate how mitochondrial respiration of PBMCs is affected in postmenopausal EBC patients by adjuvant chemotherapy administered with or without radiotherapy.

## Materials and methods

The clinical study was conducted according to the Declaration of Helsinki and approved by the Ethics Committee of The Capital Region, Denmark (Project number H-18016600 and 67762). Informed consent was obtained from all patients involved in the study.

### Clinical study

The clinical study has been described previously^14^. Briefly, postmenopausal EBC patients eligible for adjuvant chemotherapy were recruited to the clinical trial named *Healthy Living after Breast Cancer* (NCT03784651) at the Department of Oncology, Rigshospitalet, Copenhagen, Denmark. All postmenopausal patients included were aged 50-70 years, diagnosed with stage I-III EBC, and had not yet started chemotherapy or local radiotherapy at the time of recruitment. Exclusion criteria were preexisting endocrine or metabolic disease (e.g., type 2 diabetes, osteoporosis, or thyroid disease) and malignancy prior to the current breast cancer diagnosis. Patients included in the clinical trial are continuously examined every six months in the first two years and afterwards yearly for five years in this observational, longitudinal follow-up study. Data presented in this article include 23 EBC patients examined before adjuvant chemotherapy (pre treatment), as well as after completion of chemotherapy with or without radiotherapy (post treatment). Tumor characteristics and treatment regimens of the patients are summarized in Supplemental Table 1. Most patients received a combination of paclitaxel, cyclophosphamide, and epirubicin as adjuvant chemotherapy (78%), with an average time between the first and last treatment of 112 days (ranging from 85 to 163 days). Seven of the patients received neoadjuvant chemotherapy and eighteen patients received local radiotherapy. In this paper, adjuvant therapy will refer to all patients receiving chemotherapy with or without radiotherapy. Post-treatment visits were scheduled as close to therapy completion as possible, typically within three months. Patients with estrogen-positive disease had initiated endocrine therapy (aromatase inhibitors) at their post-visit (n = 20). Anthropometric measurements (e.g., height, weight, and body mass index) and blood sampling of the patients were performed at the Department of Endocrinology and Metabolism, Rigshospitalet^14^. Tumor characteristics, surgery, and antineoplastic treatment regimens were documented at the Department of Oncology, Rigshospitalet (Supplemental Table 1) and analysis of blood samples (fasting glucose, fasting insulin, HbA1c, total cholesterol, HDL cholesterol, LDL cholesterol, triglyceride, C peptide, hemoglobin, leukocytes) were performed at the Department of Clinical Biochemistry, Rigshospitalet. Initial data on 38 patients included in the *Healthy Living after Breast Cancer* study are published^14^. Here, we present data on 23 patients, 9 of whom were included in the publication by Buch-Larsen et. al.^14^.

### Peripheral blood mononuclear cell isolation

Venous blood (9 or 15 mL) was sampled in K_2_EDTA tubes, diluted in PBS to a final volume of 30 mL, carefully layered on top of 15 mL Lymphoprep™ density gradient medium (StemCell Technologies Inc., Vancouver, Canada), and centrifuged at 800 x g for 15 min (RT, acceleration 9, deceleration 2) to separate PBMCs from granulocytes and erythrocytes. PBMCs were gently collected with a Pasteur pipette and transferred to a separate tube. Isolated PBMCs were washed with PBS and centrifuged at 433 x g for 5 min (RT, acceleration 9, deceleration 9). The supernatant was discarded and the PBMC pellet was re-suspended in 1 mL of Mitochondrial Respiration media 05 (MiR05, content described below). The amount of PBMCs was quantified by the automated cell counter NucleoCounter® NC-3000 using ChemoMetec NucleoView NC-3000 software (ChemoMetec, Lillerød, Denmark).

### Polarographic measurement of oxygen consumption

Respiration was measured by high-resolution respirometry (HRR) using Oxygraph-O2k instruments (Oroboros Instruments, Innsbruck, Austria) at 37°C and 750 rpm stirring. Oxygen consumption over time and the derivative oxygen flux (O_2_ flux) was recorded by DatLab 6.1.0.7 software (Oroboros Instruments). Instrumental background calibration was routinely performed according to the manufacturer’s guidelines. All experiments were performed in 2 mL chambers containing MiR05 (110 mM sucrose, 60 mM potassium lactobionate, 0.5 mM EGTA, 3 mM MgCl_2_ · 6H_2_O, 20 mM taurine, 10 mM KH_2_PO_4_, 20 mM HEPES, 1 g/L BSA, pH 7.1). Oxygen calibration at air saturation was performed prior to each experiment. Measurements of O_2_ flux per million PBMCs were based on the current barometric pressure, the constantly measured oxygen concentration in the chamber, and the oxygen solubility factor of MiR05 (0.92)^44^. Experiments were performed at an oxygen concentration ranging from 0 to 200 μM. Oxygen concentration and O_2_ flux were measured by polarographic oxygen sensors over time, determining the oxygen consumption rate per million PBMCs. Based on PBMC quantification results, a cell suspension volume corresponding to 1.5 million PBMCs was determined. This volume of MiR05 was removed from the Oxygraph-O2k chambers, and the same volume of cell suspension was added. Substrates and inhibitors were added to the Oxygraph-O2k chambers to distinguish oxygen consumption attributed to different mitochondrial complexes and respiratory states. Two protocols were applied, and measurements were performed in duplicate as followed:

#### Intact PBMCs

Endogenous routine respiration (no substrate addition); Oligomycin (4 μg/mL, to estimate proton leak over the inner mitochondrial membrane (incl. non-mitochondrial respiration)); FCCP titration (0.25 μM, titration until the decline of O_2_ flux, to estimate the maximal capacity of the electron transport system (ETS)). This protocol examined the cellular respiration of intact PBMCs with endogenous substrates available. Oligomycin inhibits ATP synthase and FCCP uncouples OXPHOS from the electron transport chain by disturbing the electrochemical gradient across the inner mitochondrial membrane^45^.

#### Permeabilized PBMCs

Endogenous routine respiration (no substrate addition); Digitonin (5 μg/mL DMSO, to permeabilize the PBMCs); Malate (2 mM), Glutamate (10 mM), and Pyruvate (5 mM, to estimate leak respiration with the presence of complex I-linked substrates (LEAK_CI_)); ADP (5 mM) and MgCl_2_ (3 mM, to estimate complex I-linked respiration (CI_*P*_)); Cytochrome C (0.01 mM, to evaluate the integrity of the outer mitochondrial membrane); Succinate (10 mM, to estimate complex I+II-linked respiration (CI+CII_*P*_)); FCCP titration (0.25 μM, titration until the decline of O_2_ flux, to estimate maximal ETS capacity). This protocol examined O_2_ flux attributed to different mitochondrial complexes (CI_*P*_ and CI+CII_*P*_) and specific respiratory states (LEAK_CI_ and ETS) of PBMCs with saturating concentrations of substrates and inhibitors. Digitonin selectively permeabilizes the plasma membrane. Malate, glutamate, and pyruvate are complex I-linked substrates, while succinate is a complex II-linked substrate.

### Statistical analysis

Statistical analyses were performed using RStudio (v.1.3.1093). Before statistical analyses were conducted, normal distribution of data was assessed by QQ-plots and Shapiro-Wilk tests. Moreover, variance equality was assessed by F tests and homoscedasticity plots of residuals. If assumptions for statistical tests were not met, data were log-transformed. This is stated in figure legends. To statistically test changes in clinical variables paired Student’s *t*-tests were performed comparing parametric data pre and post adjuvant therapy. Wilcoxon signed-rank tests were used for non-parametric data. Changes in PBMC respiration following adjuvant therapy were analyzed by a linear mixed effect model with time (pre versus post adjuvant therapy) as a fixed effect and patient IDs as a random effect. Changes in respiratory capacities of intact PBMCs and respiratory control ratios of permeabilized PBMCs after compared to before adjuvant therapy were tested by paired Student’s *t*-tests. Specifications of statistical tests performed on presented data are stated in figure and table legends. A *p*-value of < 0.05 was considered statistically significant.

## Results

### Changes to leukocyte count and lipid profile in EBC patients after adjuvant therapy

Anthropometric and biochemical data of the EBC patients pre and post adjuvant therapy are shown in Table 1. Average age of the patients was 59 years. The patients did not change weight following adjuvant therapy and there were no significant changes in fasting glucose, fasting insulin, C peptide, or HbA1c levels. When assessing the fasting lipid profile of the EBC patients after compared to before adjuvant therapy, we found significantly decreased HDL cholesterol levels and increased LDL cholesterol and triglyceride levels (Table 1). Moreover, the hemoglobin level and leukocyte count were significantly decreased after completion of adjuvant therapy. These results are in accordance with previously published results from this clinical study^14^. Data on differential blood count is not available from this study cohort.

**Table 1.** Clinical data of postmenopausal EBC patients before and after adjuvant therapy.

|  | Pre adjuvant therapy<br>(n = 23) | Post adjuvant therapy<br>(n = 23) | Paired *t*-test<br>*p*-value |
| --- | --- | --- | --- |
| Age (years) | 59 ± 5 | N/A | N/A |
| Weight (kg) | 75 ± 12 | 74 ± 12 | *p* = 0.43 |
| BMI (kg/m<sup>2</sup>) | 27 ± 4 | 27 ± 4 | *p* = 0.68 |
| Fasting glucose (mmol/L) <sup>1</sup> | 5.9 ± 1.0 | 5.8 ± 1.4 | *p* = 0.49 |
| Fasting insulin (pmol/L) <sup>1</sup> | 76 (62, 116) | 89 (69, 129) | *p* = 0.36 |
| C peptide (pmol/L) <sup>1</sup> | 780 (640, 1090) | 947.0 (747, 947) | *p* = 0.49 |
| HbA1c (mmol/mol) | 37.0 (35.0, 40.0) | 37.0 (36.0, 39.0) | *p* = 0.82 |
| Total cholesterol (mmol/L) <sup>1</sup> | 5.9 ± 1.1 | 6.1 ± 0.9 | *p* = 0.12 |
| LDL cholesterol (mmol/L) <sup>1</sup> | 3.8 ± 0.9 | 4.1 ± 0.8 | *p* = 0.02 |
| HDL cholesterol (mmol/L) <sup>1</sup> | 1.8 ± 0.4 | 1.6 ± 0.2 | *p* = 0.004 |
| Triglyceride (mmol/L) <sup>1</sup> | 1.3 ± 0.4 | 1.6 ± 0.8 | *p* = 0.01 |
| Hemoglobin (mmol/L) <sup>2</sup> | 8.2 ± 0.8 | 7.7 ± 0.5 | *p* = 0.0003 |
| Leukocytes (10<sup>9</sup>/L) <sup>2</sup> | 5.7 ± 1.1 | 4.7 ± 1.1 | *p* = 1 x 10<sup>-4</sup> |

Data are presented as mean ± SD for normally distributed variables and median (25-75% interquartile range) for variables that were not normally distributed. BMI: Body Mass Index; HbA1c: Hemoglobin A1c; LDL: Low Density Lipoprotein; HDL: High Density Lipoprotein; N/A: Not applicable. <sup>1</sup> n = 21 for indicated parameters post adjuvant therapy as two patients were not assessed in a fasted state. <sup>2</sup> n = 22. Statistically significant changes were identified by paired Student's *t*-test for parametric data and Wilcoxon signed-rank test for non-parametric data.

### Mitochondrial respiration in PBMCs from EBC patients is increased after adjuvant therapy

Noticeably, the mitochondrial respiratory capacity of various states in intact PBMCs from the EBC patients after treatment was significantly increased compared to before adjuvant therapy (*p* = 6 × 10^−5^, Figure 1). We found no significant interaction between adjuvant therapy and respiratory states, indicating that there was no significant difference in how endogenous respiration, proton leak, and maximal respiratory capacity of the ETS responded to the course of therapy.

**Figure 1.**
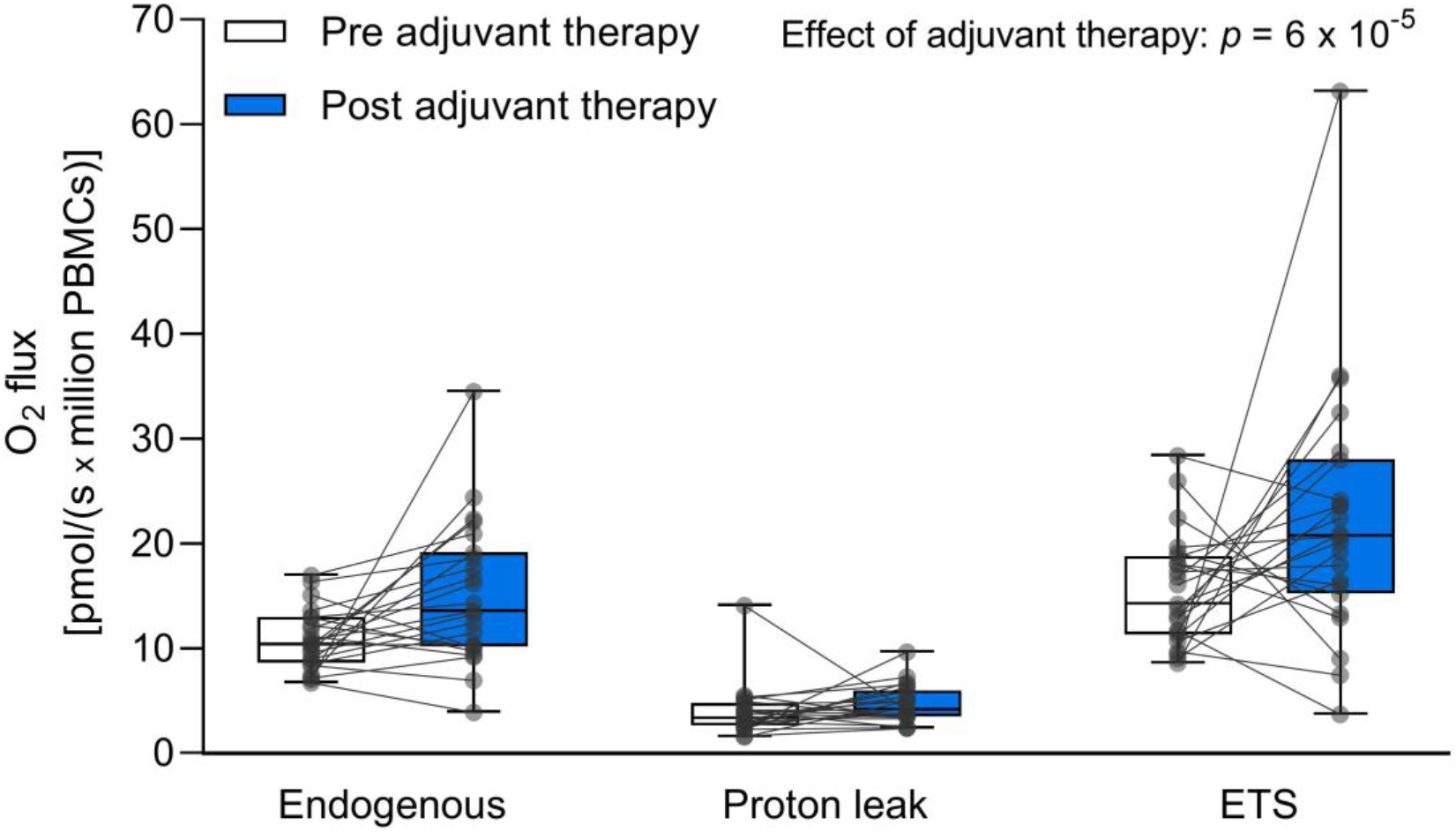
Respiration of intact PBMCs from EBC patients pre and post adjuvant therapy. Boxplots indicate medians, interquartile range (25^th^ to 75^th^ percentile) and minimum to maximum (whiskers) O_2_ flux. All participants are shown as individual data points. White boxes: Pre adjuvant therapy, n = 23. Blue boxes: Post adjuvant therapy, n = 23. Endogenous: Endogenous routine respiration (no substrates or inhibitors added). Proton leak: Oxygen consumed due to proton leak over the inner mitochondrial membrane including a non-mitochondrial respiration contribution. ETS: Maximal capacity of the electron transport system (uncoupled state). Linear mixed effects model analysis identified adjuvant therapy as the main effect on the respiratory states in intact PBMCs. Data were log-transformed prior to the statistical analysis.

To evaluate if the increased respiration of intact PBMCs was driven by specific mitochondrial complexes or respiratory states, HRR analysis of permeabilized PBMCs from the EBC patients was performed. Respiration of permeabilized PBMCs (O_2_ flux per million cells) was significantly increased after adjuvant therapy compared to before (*p* = 1 × 10^−7^, Figure 2). There was no significant interaction between adjuvant therapy and the different mitochondrial complexes and respiratory states, indicating that the observed effect of adjuvant therapy on O_2_ flux was not significantly different for specific complexes of the electron transport chain or respiratory states.

**Figure 2.**
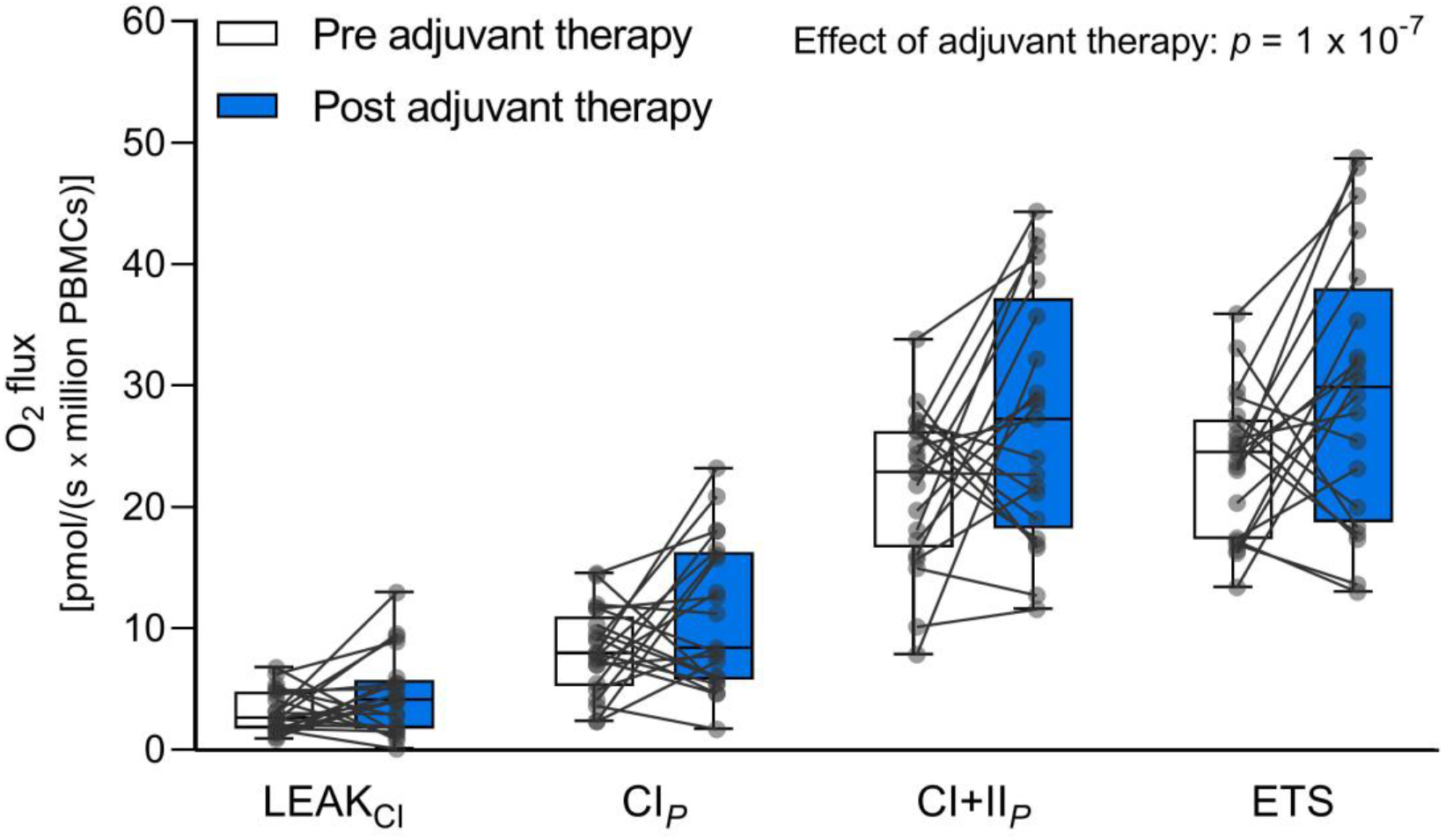
Mitochondrial respiration of permeabilized PBMCs from EBC patients pre and post adjuvant therapy. Data are represented by boxplots indicating medians, interquartile range (25^th^ to 75^th^ percentile) and minimum to maximum (whiskers) O_2_ flux. All participants are shown as individual data points. White boxes: Pre adjuvant therapy, n = 21. Blue boxes: Post adjuvant therapy, n = 21. LEAK_CI_: Leak respiration with the presence of complex I-linked substrates. CI_*P*_: Complex I-linked respiration. CI+II_*P*_: Complex I+II-linked respiration. ETS: Maximal capacity of the electron transport system, n = 20. Linear mixed effect model analysis identified adjuvant therapy as the main effect on the O_2_ flux of mitochondrial complexes and respiratory states of permeabilized PBMCs.

When assessing the changes in O_2_ flux for specific mitochondrial complexes and respiratory states separately, we found that the O_2_ consumption was increased for all respiratory parameters after adjuvant therapy, ranging from 17 % (Proton leak) to 43 % (ETS; Table 2). When evaluating the respiratory capacities of intact PBMCs and respiratory control ratios of permeabilized PBMCs (Table 3), we found the O_2_ consumption used for oxidative phosphorylation of ADP into ATP in intact PBMCs (ATP-linked OXPHOS) to be significantly increased after adjuvant therapy compared to before (*p* = 0.008).

**Table 2.** Percentage change in O_2_ flux from pre to post adjuvant therapy for specific mitochondrial complexes and respiratory states of PBMCs from EBC patients.

|  | Percentage change in O<sub>2</sub> flux<br>[pmol/(s x million PBMCs)] |
| --- | --- |
| Intact PBMCs |  |
| Endogenous | + 40 % |
| Proton leak | + 17 % |
| ETS | + 43 % |
| Permeabilized PBMCs |  |
| Endogenous | + 38 % |
| LEAK<sub>CI</sub> | + 42 % |
| CI<sub>P</sub> | + 35 % |
| CI+II<sub>P</sub> | + 26 % |
| ETS | + 25 % |

In the HRR protocol for permeabilized PBMCs, endogenous respiration was assessed before the addition of digitonin.

**Table 3.** Respiratory capacities of intact PBMCs and respiratory control ratios of permeabilized PBMCs from EBC patients pre and post adjuvant therapy.

|  | Pre adjuvant therapy | Post adjuvant therapy | Change (%) | Paired *t*-test *p*-value |
| --- | --- | --- | --- | --- |
| Intact PBMCs |  |  |  |  |
| ATP-linked OXPHOS | 6.85 ± 2.13 | 10.48 ± 5.35 | +53% | 0.008 |
| Reserve capacity | 4.73 ± 3.21 | 7.12 ± 6.23 | +51% | 0.19 |
| Permeabilized PBMCs |  |  |  |  |
| LEAK<sub>CI</sub> / OXPHOS | 0.15 ± 0.10 | 0.15 ± 0.09 | 0% | 0.96 |
| CI<sub>P</sub> / CI+II<sub>P</sub> | 0.37 ± 0.12 | 0.38 ± 0.11 | +3% | 0.76 |
| OXPHOS / ETS | 0.90 ± 0.12 | 0.94 ± 0.07 | +4% | 0.28 |
| R / ETS | 0.49 ± 0.08 | 0.53 ± 0.12 | +8% | 0.24 |

Data are presented as mean + SD of respiratory capacities of intact PBMCs (n = 23) and respiratory control ratios of permeabilized PBMCs (n = 21). ATP-linked OXPHOS: Oxygen consumption used for oxidative phosphorylation (Endogenous – Leak). Reserve capacity: Respiratory reserve capacity from endogenous routine respiration to theoretical maximal capacity of the ETS (uncoupled state; ETS – Endogenous). LEAK<sub>CI</sub>/OXPHOS: Proportion of OXPHOS attributed to leak respiration with the presence of complex I-linked substrates. CI<sub>P</sub>/CI+II<sub>P</sub>: Contribution of complex I to OXPHOS at fully saturated substrate concentrations. OXPHOS/ETS: Part of the maximal respiratory capacity of the ETS which OXPHOS uptakes, n = 20. R/ETS: Respiratory reserve capacity from routine respiration (R) to theoretical maximal capacity of the ETS, n = 16. Statistically significant changes were identified by paired Student's *t*-test.

Next, we investigated the impact of different treatment regimens on the endogenous respiration of intact PBMCs. EBC patients were stratified into two groups according to if they received radiotherapy (chemotherapy and radiation, n = 18) or not (chemotherapy only, n = 5). Also, EBC patients who did not receive the standard chemotherapy drug combination (cyclophosphamide, epirubicin, and paclitaxel) were highlighted. As shown in Figure 3, the significant increase in PBMC respiration after adjuvant therapy does not seem noticeably distinctive for different chemotherapeutic drug combinations. The increased endogenous respiration seems to be less prominent in patients receiving chemotherapy alone (n = 5) compared to patients receiving combined chemo- and radiotherapy (n = 18). However, this has not been statistically tested as we recognize that the sample size in the stratified groups is relatively small, rendering the interpretation of these effects challenging.

**Figure 3.**
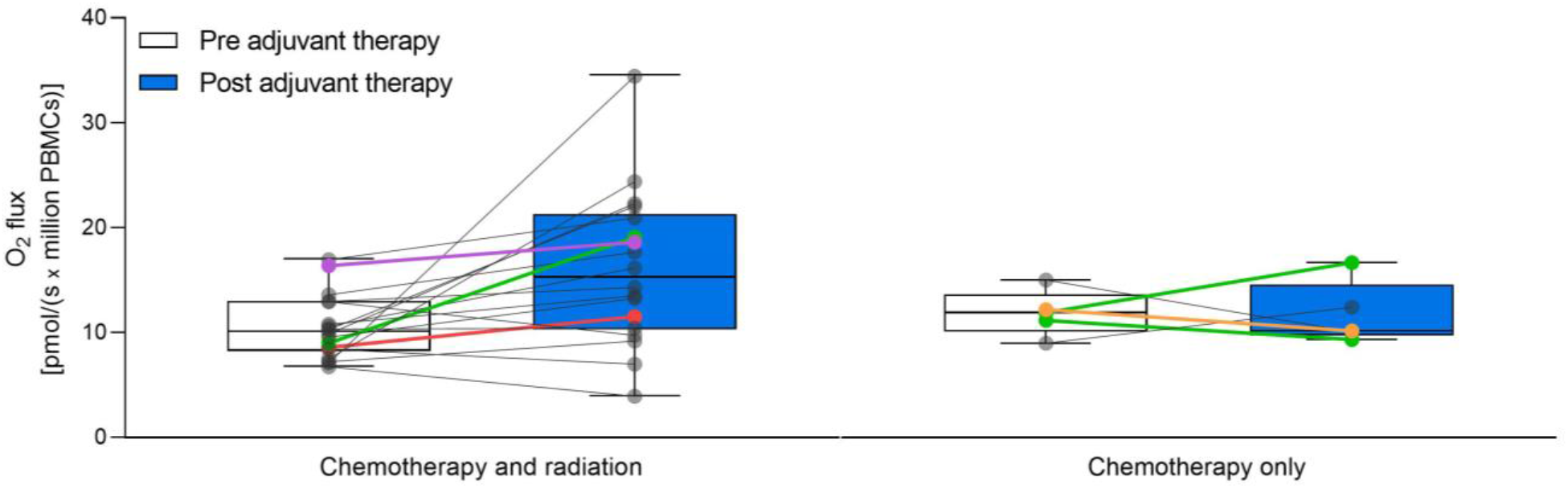
Endogenous respiration of intact PBMCs from EBC patients pre and post adjuvant therapy. Endogenous oxygen consumption measured as O_2_ flux per million intact PBMCs from EBC patients pre and post adjuvant therapy is presented by boxplots indicating medians, interquartile range (25^th^ to 75^th^ percentile) and minimum to maximum (whiskers) O_2_ flux. All participants are shown as individual data points. EBC patients are stratified according to treatment regimen, chemo- and radiotherapy (n = 18) or chemotherapy only (n = 5). Color of the data points indicates the chemotherapeutic drug combination used to treat the specific patient. Grey: Cyclophosphamide, epirubicin, and paclitaxel, n = 15 (chemo- and radiotherapy) and 2 (chemotherapy only). Green: Paclitaxel, n = 1 (chemo- and radiotherapy) and 2 (chemotherapy only). Red: Cyclophosphamide, epirubicin, paclitaxel, and capecitabine, n = 1. Purple: Cyclophosphamide and docetaxel, n = 1. Orange: Cyclophosphamide and epirubicin, n = 1. White boxes: Pre adjuvant therapy. Blue boxes: Post adjuvant therapy.

### No associations between PBMC respiratory capacity and metabolic characteristics

Lastly, we investigated if endogenous PBMC respiration, proton leak or ETS capacity showed any association with age, BMI or the levels of fasting LDL cholesterol, HDL cholesterol, triglycerides or glucose levels in the EBC patients before adjuvant treatments, which has previously been observed in healthy human subjects^46^. We found no significant association between endogenous respiration, proton leak or maximal ETS capacity of intact PBMCs and any of the clinical parameters (Supplemental Table 2).

## Discussion

In this study, we investigated how adjuvant chemotherapy with or without radiotherapy affected the mitochondrial respiration of PBMCs in postmenopausal EBC patients. Interestingly, we found that respiration of intact PBMCs and respiration attributed to specific mitochondrial complexes and respiratory states of permeabilized PBMCs were significantly increased after adjuvant chemo- and radiotherapy treatment compared to before. Thus, our data suggest that PBMCs in EBC patients require more oxygen to support cellular functions after therapy completion. The impact on specific mitochondrial complexes and respiratory states was not significantly different, indicating a generally increased oxygen consumption of PBMCs post adjuvant therapy which was not driven by specific components of the mitochondria. However, ATP-linked OXPHOS in intact PBMCs was significantly increased and since proton leak was increased to a lower extent than other respiratory states, this could together indicate a better coupling of the electron transport chain to OXPHOS in the PBMCs after adjuvant therapy. The number of patients included in this study limits evaluation of the impact of different treatment regimens on PBMC respiration, but the increase was more pronounced in patients receiving both chemo- and radiotherapy (78% of our patients).

The population of mononuclear cells in peripheral blood is heterogeneous and consists of T cells, B cells, NK cells, monocytes, and dendritic cells in varying amounts^47^. Utilization of OXPHOS for energy production is different in the different cell types^48^, thus alterations in proportions of cell types after versus before adjuvant therapy could contribute to the observed increase in oxygen consumption. Previously reported effects of chemo- and radiotherapy on different types of leukocytes in peripheral blood from breast cancer patients are not consistent. Some^49–53^ but not all^54^ studies report significant reductions in total lymphocytes and/or total number of T cells with a persistent decrease in T helper (CD4+) cells in breast cancer patients after chemo- and/or radiotherapy.

T cells comprise 45-70% of isolated mononuclear cells from human peripheral blood^55^. T cells that primarily rely on mitochondrial OXPHOS for ATP generation are resting naïve T cells, antigen-specific memory T cells, and FoxP3+ regulatory T cells, whereas proliferative helper (CD4+) and cytotoxic (CD8+) effector T cells utilize glycolysis to a higher extent^56–59^. Energy metabolic profiling of B lymphocyte subsets and NK cells is less well described^48^. Although speculative, the increased mitochondrial respiration of PBMCs post adjuvant therapy found in this study could indicate a shift in the subpopulation of T cells from effector T cells to naïve T cells, memory T cells, and/or regulatory T cells with higher OXPHOS utilization. Such a shift could be caused by different sensitivity, turnover, and/or recovery of PBMC subpopulations in response to adjuvant treatments^24,49,51^. Unfortunately, we do not have differential blood counts from the EBC patients included in this study.

It is well known that repeated cycles of chemo- and radiotherapy can induce damage to the hematopoietic system^61,62^. Previous research have shown that hematopoietic progenitor cells (HPCs) can be depleted by chemo- and/or radiotherapy after which hematopoiesis primarily will be restored by hematopoietic stem cells (HSCs)^61^. Adjuvant therapy administered to our patients might have eliminated existing HPCs and concurrently affected the bone marrow microenvironment and thereby the generation of new blood cells. In case the proportion of newly differentiated PBMCs is increased after adjuvant therapy, OXPHOS utilization might be increased due to their naïve phenotype. However, proportion of PBMC subpopulations during and after completion of adjuvant chemotherapy administration of averagely 112 days might also vary because of e generally fast leukocyte cell turnover. Importantly, because of the predominantly short lifespan of white blood cells^60^, the high degree of PBMC regeneration compared to more long-lived cells in tissues could explain why we see increased respiration in PBMCs as opposed to impaired mitochondrial function in e.g., murine skeletal myocytes or cardiomyocytes after chemotherapy exposure^27,28,30–32^. Therefore, translation of the effect of adjuvant therapy on long-lived cells to short-lived cells such as PBMCs must be done with caution.

Besides the possible explanations discussed, the increased respiration can also be caused by an increased number of mitochondria in the PBMCs. Mitochondrial content loss has been shown in skeletal myocytes from EBC patients after (neo)adjuvant chemotherapy^35^, but it has not been addressed in PBMCs. CD8+ memory T cells have been shown to have enhanced mitochondrial content compared to CD8+ effector T cells^59^, supporting a possible shift in immune cell subpopulations during adjuvant therapy towards increased OXPHOS utilization as described. Previous studies have also shown that DNA damage induced by ionizing radiation cause up-regulation of mitochondrial biogenesis in human glioblastoma cells^63^. Thus, the therapy-induced increase in PBMC respiration in the present study might also be related to the effects of adjuvant therapy on the mitochondrial amount in PBMCs. In summary, the increased respiratory O_2_ flux in PBMCs from EBC patients after adjuvant therapy might be attributed to an altered cell population with a higher proportion of cell types relying on OXPHOS rather than glycolysis, a direct effect of adjuvant therapy on the bone marrow leading to altered hematopoiesis, or an increased mitochondrial content in the PBMCs.

Metabolic factors may influence the vasculature and thereby the microenvironment of peripheral blood cells. The EBC patients included in this study exhibited decreased HDL cholesterol levels and increased triglyceride and LDL cholesterol levels post versus pre adjuvant therapy, which is in accordance with the findings of other studies^9,14^. Importantly, DeConne et al. found a negative correlation between LDL cholesterol levels and maximal respiration as well as spare respiratory capacity in PBMCs from healthy adults, suggesting that there may be a link between blood cholesterol and mitochondrial respiration of PBMCs in humans^46^. However, such associations were not evident from our data.

It remains unknown what causes the PBMCs to have an increased oxygen demand after adjuvant therapy completion and whether the changes are persistent or will normalize during the first year after chemotherapy completion. This study is to our knowledge the first to investigate how energy metabolism in PBMCs is affected with adjuvant chemo- and radiotherapy treatment of postmenopausal EBC patients. We encourage more studies to evaluate the effects of adjuvant therapy on the metabolic as well as immunologic state of healthy cells and tissues from the large and growing population of breast cancer patients.

## Conclusion

This study shows that postmenopausal EBC patients treated with chemotherapy with or without radiotherapy exhibit increased mitochondrial respiration in circulating peripheral blood mononuclear cells after therapy completion compared to before. The increased oxygen demand of PBMCs after adjuvant chemo- and radiotherapy could be explained by a shift in PBMC subpopulation proportions towards cells favoring OXPHOS over glycolysis, an increased mitochondrial content in PBMCs, and/or therapy-induced alterations in the hematopoietic system or the vascular microenvironment wherein the PBMCs reside. However, these mechanisms need further investigation in larger observational studies.

## Abbreviations

CI_*P*_: Complex I-linked respiration
CI+CII_*P*_: Complex I+II-linked respiration
EBC: Early breast cancer
ER: Estrogen receptor
ETS: Electron transport system
FACS: Fluorescence-activated cell sorting
FCCP: Carbonylcyanide *p*-trifluoromethoxyphenylhydrazone
HbA1c: Hemoglobin A1c
HDL: High density lipoprotein
HER2: Human Epidermal Growth Factor Receptor 2
HPC: Hematopoietic progenitor cell
HRR: High-resolution respirometry
HSC: Hematopoietic stem cell
LDL: Low density lipoprotein
LEAK_CI_: Leak respiration through complex I
MiR05: Mitochondrial respiration media 05
OXPHOS: Oxidative phosphorylation
PAM50: Prediction Analysis of Microarray 50
PBMC: Peripheral blood mononuclear cell
R: Routine respiration.

## Acknowledgement

We thank all participants who volunteered to enroll in the study. We also thank Malan Egholm, Rigshospitalet, and Regitze Kraunsøe, University of Copenhagen, for excellent technical assistance. *Financial support:* This study was financially supported by the Danish Diabetes Academy (grant number NNF17SA0031406), Fru Astrid Thaysens Legat for Lægevidenskabelig Grundforskning, Dagmar Marshalls Fond, Svend Andersen Fonden and Aase og Ejnar Danielsens Fond. *Clinical Trial Information:* NCT03784651 (registered 24 December 2018).

