## Supplemental data for "Peripheral blood mononuclear cells exhibit increased mitochondrial respiration after adjuvant chemo- and radiotherapy for early breast cancer"

**Supplemental Table 1** | Tumor characteristics and treatment regimens of EBC patients.

| Number of patients | n = 23 |
|---|---|
| Tumor stage | |
| I | 5 (22 %) |
| II | 14 (61 %) |
| III | 4 (17 %) |
| Histology | |
| Invasive Ductal Carcinoma | 21 (91 %) |
| Invasive Luminal/Lobular Carcinoma | 2 (9 %) |
| Laterality | |
| Right | 11 (48 %) |
| Left | 10 (43 %) |
| Bilateral | 2 (9 %) |
| Surgery type | |
| Mastectomy | 7 (30 %) |
| Lumpectomy | 16 (70 %) |
| Lymph node involvement | |
| 0 | 7 (30 %) |
| 1-3 | 16 (70 %) |
| 4+ | 0 (0 %) |
| ER status | |
| Positive | 20 (87 %) |
| Negative | 3 (13 %) |
| HER2 status | |
| Positive | 9 (39 %) |
| Negative | 14 (61 %) |
| PAM50 [1] | |
| Luminal A | 3 (16 %) |
| Luminal B | 9 (47 %) |
| Luminal C | 3 (16 %) |
| HER2 enriched | 0 (0 %) |
| Basal like | 0 (0 %) |
| Chemotherapy | |
| Cyclophosphamide | 20 (87 %) |
| Epirubicin | 19 (83 %) |
| Docetaxel | 1 (4 %) |
| Paclitaxel | 21 (91 %) |
| Capecitabine | 1 (4 %) |
| Radiotherapy | |
| Yes | 18 (78 %) |
| No | 5 (22 %) |
| Number of radiation cycles | 15 (67%), 25 (28%) or 30 (5%) |
| Endocrine treatment (aromatase inhibitors) | |
| Yes | 20 (87 %) |
| No | 3 (13 %) |

Data are presented as number of patients and percentage of the cohort specified in parenthesis. Number of radiation cycles are presented as median and 25-75% interquartile range. ER: Estrogen Receptor; HER2: Human Epidermal Growth Factor Receptor 2; PAM50: Prediction Analysis of Microarray 50; [1]n=19.

**Supplemental Table 2** | Correlations between intact PBMC respirometry and clinical characteristics of postmenopausal EBC patients before adjuvant therapy.

| | **Endogenous routine** (n = 23) | **Proton leak** (n = 23) | **ETS** (n = 23) |
|---|---|---|---|
| Age (years) | r=0,1883 (-0,2549 to 0,5661) | r=0,2997 (-0,1410 to 0,6413) | r=-0,02626 (-0,4442 to 0,4011) |
| BMI (kg/m$^2$) | r=0,1354 (-0,3049 to 0,5281) | r=-0,1448 (-0,5350 to 0,2962) | r=0,1265 (-0,3131 to 0,5215) |
| Fasting glucose (mmol/L) | r=0,2793 (-0,1628 to 0,6281) | r=-0,03234 (-0,4491 to 0,3960) | r=0,2179 (-0,2258 to 0,5868) |
| Fasting insulin (pmol/L) | r=0,2625 (-0,1804 to 0,6169) | r=0,1097 (-0,3284 to 0,5090) | r=0,2111 (-0,2326 to 0,5820) |
| Total cholesterol (mmol/L) | r=0,03373 (-0,3948 to 0,4502) | r=-0,01045 (-0,4314 to 0,4143) | r=-0,05456 (-0,4667 to 0,3770) |
| LDL cholesterol (mmol/L) | r=-0,08803 (-0,4926 to 0,3478) | r=0,1352 (-0,3052 to 0,5279) | r=-0,1726 (-0,5550 to 0,2700) |
| HDL cholesterol (mmol/L) | r=0,3027 (-0,1379 to 0,6432) | r=0,02778 (-0,3998 to 0,4454) | r=0,2604 (-0,1826 to 0,6155) |
| Triglyceride (mmol/L) | r=0,01791 (-0,4081 to 0,4375) | r=-0,03264 (-0,4493 to 0,3957) | r=-0,04776 (-0,4613 to 0,3829) |

Results are presented as Spearman correlation coefficients (r) with 95% confidence intervals. Endogenous: Endogenous routine respiration (no substrates or inhibitors added). Proton leak: Oxygen consumed due to proton leak over the inner mitochondrial membrane including a non-mitochondrial respiration contribution. ETS: Maximal capacity of the electron transport system (uncoupled state). BMI: Body Mass Index; LDL: Low Density Lipoprotein; HDL: High Density Lipoprotein. *: $p < 0.05$.
